# The secreted hypersensitive response inducing protein 1 from *Botrytis cinerea* displays non-canonical PAMP-activity

**DOI:** 10.1101/2020.12.16.423131

**Authors:** Tanja Jeblick, Thomas Leisen, Christina E. Steidele, Jonas Müller, Florian Mahler, Frederik Sommer, Sandro Keller, Ralph Hückelhoven, Matthias Hahn, David Scheuring

## Abstract

According to their lifestyle, plant pathogens are divided into biotrophic and necrotrophic organisms. While biotrophic pathogens establish a relationship with living host cells, necrotrophic pathogens rapidly kill host cells and feed on the cell debris. To this end, the necrotrophic ascomycete fungus *Botrytis cinerea* secretes large amounts of phytotoxic proteins and cell wall degrading enzymes. However, the precise role of these proteins during the infection process is unknown. Here we report on the identification and characterization of the previously unknown toxic protein hypersensitive response inducing protein 1 (Hip1), which induces plant cell death. We found the adoption of a folded protein structure to be a prerequisite for Hip1 to exert its necrosis-inducing activity in *Nicotiana benthamiana*. Localization and the induction of specific plant responses by Hip1 indicate recognition as pathogen-associated molecular pattern at the plant plasma membrane. Our results demonstrate that recognition of Hip1, even in the absence of obvious enzymatic or pore-forming activity, induces strong plant defense reactions eventually leading to plant cell death.

## Introduction

Successful plant pathogens need to overcome preformed and activated defense barriers of their hosts for infection. To overcome such barriers at the cuticle and the cell wall, small wounds or penetration sites are created by invasive pathogens, using either disruptive enzymes or physical force or both (Hématy et al., 2009). The necrotrophic fungus *Botrytis cinerea* (*Botrytis* hereafter), causal agent of the grey mould rot, has a wide host range (>1000 species) and successful infection is characterized by rapid killing of plant cells and subsequent colonization of dead tissue (Williamson et al., 2007; Dean et al., 2012). During the infection process, large amounts of cell wall degrading enzymes (CWDEs) and phytotoxic proteins are secreted, with many of them still being of unknown molecular identity.

Up to date, several cuticle and CWDEs have been functionally characterized in *Botrytis:* For instance, CutA hydrolyses cutin (van Kan et al., 1997), pectin has been shown to be degraded by the endopolygalacturonases PG1 and PG2 (Have et al., 1998; Kars et al., 2005), the xylanases Xyn11A and Xyl1 and the xyloglucanase XYG1 (Noda et al., 2010; Yang et al., 2018; Zhu et al., 2017; Brito et al., 2006) target hemicellulose, while the endo-ß-1,4-glucanase Cel5A degrades cellulose (Espino et al., 2005). Another class of proteins typically present in necrotrophic fungi are necrosis-inducing proteins (NIPs), consisting mainly of cerato-platanins and necrosis- and ethylene-inducing proteins (NEPs and NEP-like proteins, NLPs).The latter are generally suggested to function by permeabilizing the plasma membrane to facilitate pathogen penetration (Schouten et al., 2008). Indeed, NLPs from plant-pathogenic oomycetes have been shown to bind to specific membrane lipids, conducting a pore-forming activity which eventually results in cell lysis (Lenarčič et al., 2017).

In addition to their described function, several secreted *Botrytis* proteins are recognized by the plant immune system as pathogen-associated molecular patterns (PAMPs), leading to PAMP-triggered immunity (PTI). Notably, enzymatically inactive mutant proteins are still capable of inducing cell death and often short conserved amino acid (aa) sequences could be identified as putative PAMP epitopes which are recognized by pattern recognition receptors (PRRs) at the plant plasma membrane (Boutrot and Zipfel, 2017). Among them are the xylanases Xyn11A (30 aa), Xyl1 (26 aa), the cerato-platanin Spl1 (40 aa) and IEB1 (35 aa) (Noda et al., 2010; Yang et al., 2018; Frías et al., 2011; Frías et al., 2016). Spl1 and IEB1 are both without known enzymatic activity, and it has been suggested that recognition as PAMP alone might be already sufficient for induction of necrosis. Although PTI is a relatively basal plant defense response, PAMPs can induce the hypersensitive response (HR), a strong local defense response involving programmed cell death (Thomma et al., 2011; Boutrot and Zipfel, 2017). Usually the HR serves as a measure to protect plants from pathogen proliferation, but for necrotrophic pathogens plant cell death has been shown to be beneficial (Govrin and Levine, 2000).

The generation of single knockouts mutants for the majority of phytotoxic *Botrytis* proteins showed only a mild reduction in virulence (Noda et al., 2010) or no effect on pathogenicity at all (Cuesta Arenas et al., 2010; Frías et al., 2016; Zhu et al., 2017; Schouten et al., 2008). Taken also into account that the majority of the more than 200 proteins found in the *Botrytis* secretome (Espino et al., 2010; Zhu et al., 2017; Müller et al., 2018; Li et al., 2012; Frías et al., 2016) are not well characterized yet, we hypothesized that additional, yet unknown, toxic proteins might exist. In agreement, only recently two CWDEs contributing to *Botrytis* virulence have been identified in a proteomic analysis of extracellular protein (Li et al., 2020). In this study, we report on a screen of the *Botrytis* secretome for additional phytotoxic proteins, which resulted in the identification and characterization of the previously unknown secreted *hypersensitive response inducing protein 1* (Hip1). We show that the strong necrosis-inducing activity of Hip1 relies on almost the entire aa sequence and provide evidence that Hip1 has a fold similar to that of an allergen from *Alternaria alternata* (Alt a 1). Furthermore, we demonstrate that Hip1 induces early plant immune responses, suggesting a recognition as PAMP at the plant plasma membrane.

## Results

To confirm that the toxicity of the *Botrytis* secretome produced during plant infection is mainly derived from proteins (Zhu et al., 2017), it was subjected to heat treatment and digestion by proteinase K treatment prior to infiltration into *N. benthamiana* leaves (Figure S1A). After 8 h of incubation, we observed necrosis of the entire secretome-treated leaf area whereas both heat and proteinase K treatments reduced the necrotic area to ~30% and ~20%, respectively. For the identification of as yet uncharacterized phytotoxic proteins, the secretome was fractionated, using a combination of native ion exchange chromatography and size exclusion chromatography. Toxicity of the resulting fractions was assessed by infiltration in *N. benthamiana* leaves. Next, fractions that induced necrosis were subjected to MS-based protein identification and the proteins sorted by abundance and distribution in the individual fractions. Twenty-two *Botrytis* candidate genes encoding new candidate toxic proteins (Table S1), were cloned in a binary pGreen II vector, possessing a strong 35S promotor, a plant signal peptide (SP) and a hemagglutinin (HA) tag for expression control (Fig. 1A). Using *Agrobacterium*-mediated transient expression in *N. benthamiana*, the presence of all proteins could be immunologically confirmed (Fig. S2) and necrosis-inducing activity was analysed (Fig. 1B).

**Fig. 1:**
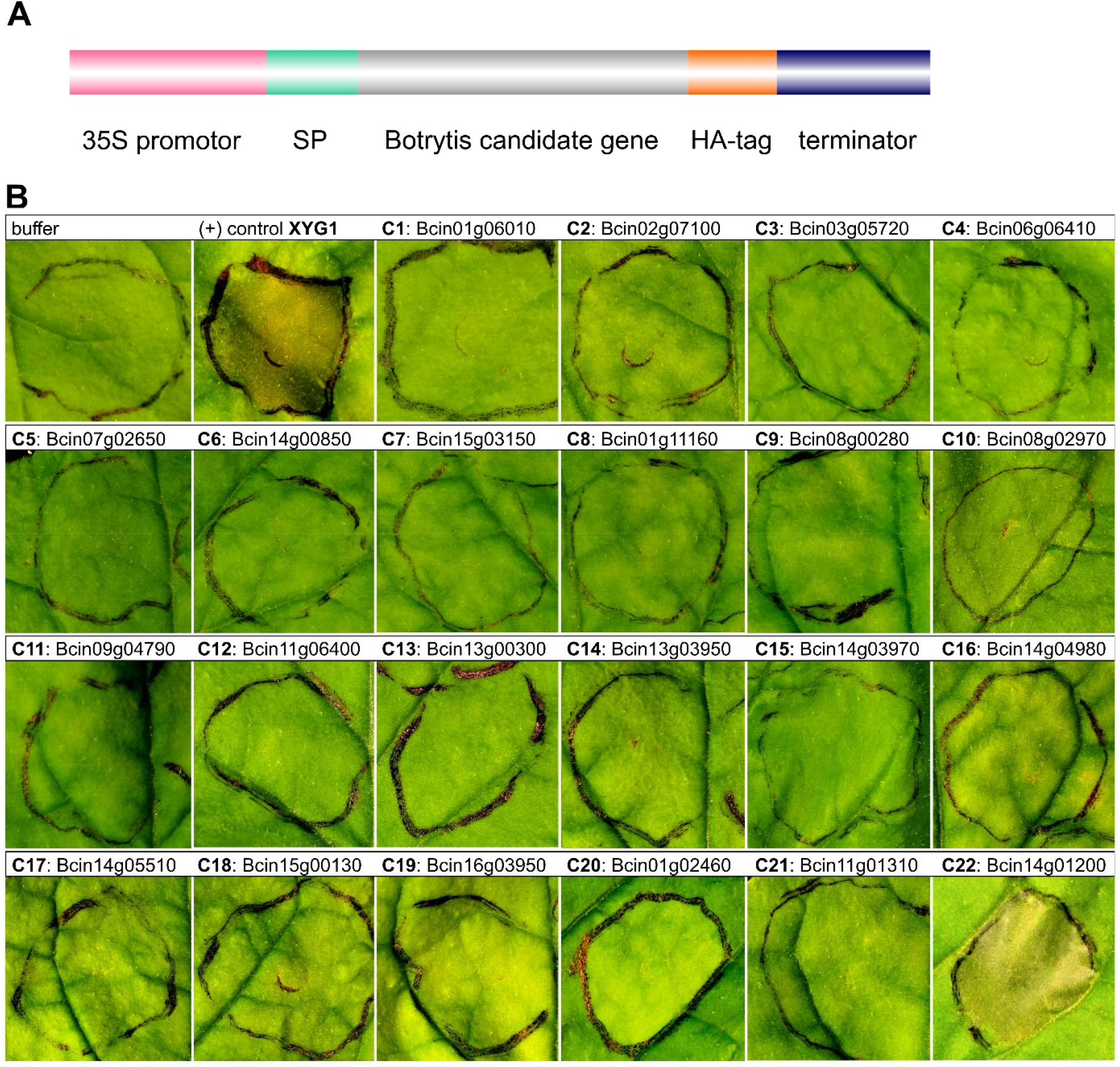
Screening for toxic *Botrytis* proteins. *Agrobacterium-mediated* transient expression using the cauliflower mosaic virus 35S promotor, a plant signal peptide (SP) and an HA tag for detection **(A)**. Symptoms on leaves of *N. benthamiana*. XYG1 (Zhu et al., 2017) was used as positive control and 22 genes encoding candidate proteins (C1-C22) were tested **(B)**. Similar to XYG1, only candidate 22 showed significant toxicity 3 days post infiltration, necrotizing the entire infiltrated leaf area.

Compared to the positive control (XYG1, Zhu et al., 2017), only candidate 22 (Bcin14g01200) showed similar necrosis-inducing activity. Previously, a homologue of candidate 22 has been described in the elm tree pathogen *Ophiostoma ulmi* as phytoalexin-inducing protein (Yang et al., 1989; Taylor et al., 2008). Lower expression levels of candidate 22 induced spot-like necrotic lesions reminiscent of the HR, and we thus named candidate 22 *hypersensitive response inducing protein 1* (Hip1). Homologs of Hip1 are found in plant pathogenic and plant-associated ascomycetous fungi, and seem to be restricted to the classes Leotiomycetes (including *Botrytis*, *Sclerotinia*, *Monilinia*), Sordariomycetes (including *Colletotrichum*, *Fusarium* and *Podospora*) and Dothideomycetes. Alignment of predicted protein sequences revealed the presence of four conserved cysteine residues (Table S2).

Since several secreted *Botrytis* proteins induce cell death in a PAMP-like manner, a search for a minimal (PAMP) cell death-inducing sequence was performed. For this, six truncated Hip1 versions, three lacking aa at the N-terminus and three lacking aa at the C-terminus were transiently expressed (Fig. 2A and 2B; Fig. S3). Only the truncation comprising aa 1-106 showed similar necrosis-inducing activity as full-length Hip1, while all other truncations were largely inactive (Fig. 2C). Examination of *Hip1* expression levels during infection revealed a strong upregulation after 48h (Fig. 2D).

**Fig. 2:**
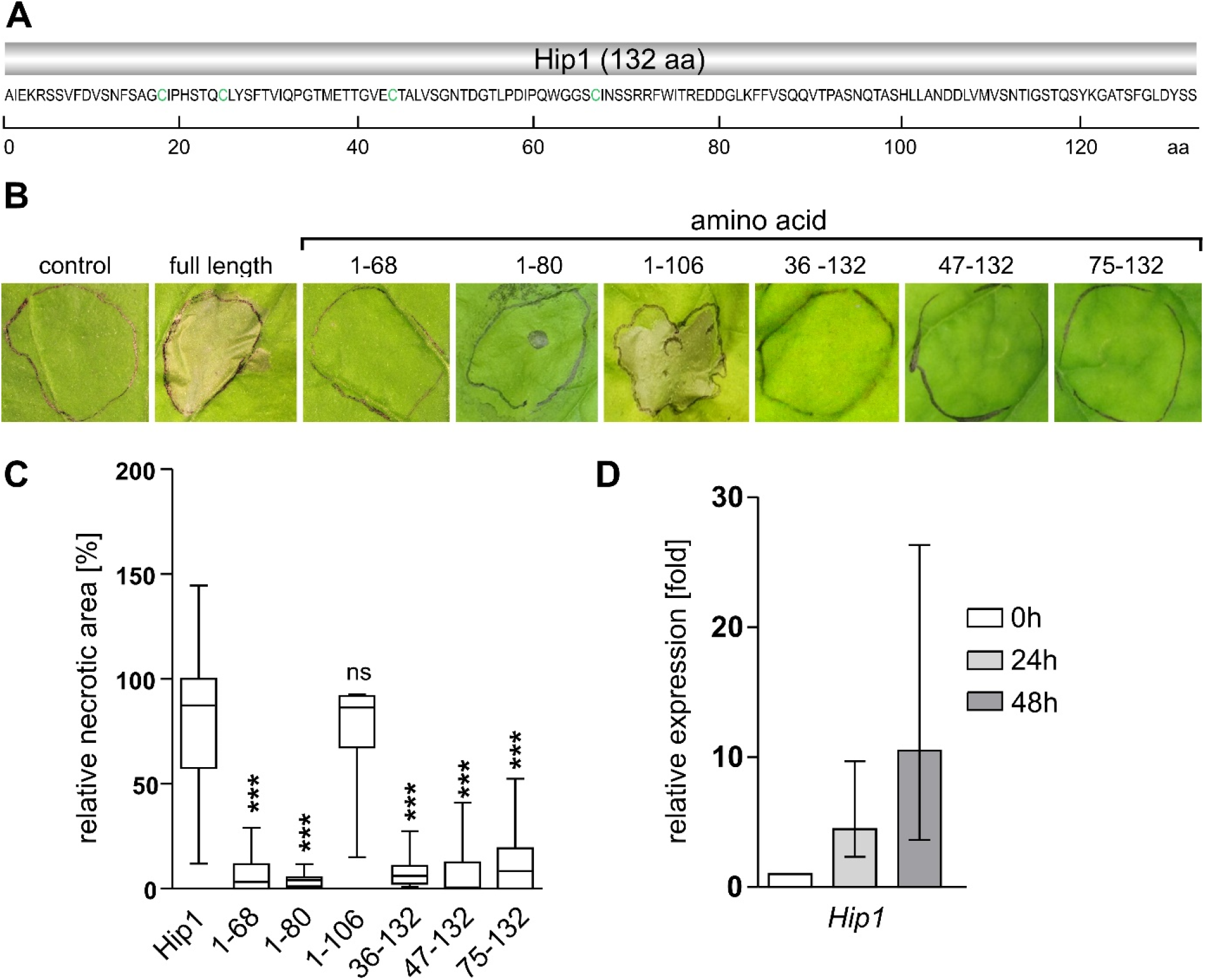
Toxicity of the hypersensitive response-inducing protein 1 (Hip1) requires the majority of the underlying amino acid sequence. Excluding the signal peptide, Hip1 is composed of 132 aa **(A)**. Conserved cysteine residues are highlighted in green. Different truncations were transiently expressed in *N. benthamiana* **(B)**. All but the truncation comprising of aa 1-106 lost toxic activity **(C)**. One-way ANOVA followed by Tukey’s post hoc test with Hip1 as control. *p-value:* *** p<0.001; n>8. qRT-PCR to assess *Hip1* expression upon infection by *Botrytis*. Values are normalized to the 0h control **(D)**.

To confirm the results from transient expression, recombinant protein expression in *E. coli* and subsequent purification was employed (Fig. S4A and S4B). Starting at 0.5 μM concentration, Hip1 showed necrosis-inducing activity 30 h post infiltration in *N. benthamiana* (Fig. 3A).

**Fig. 3:**
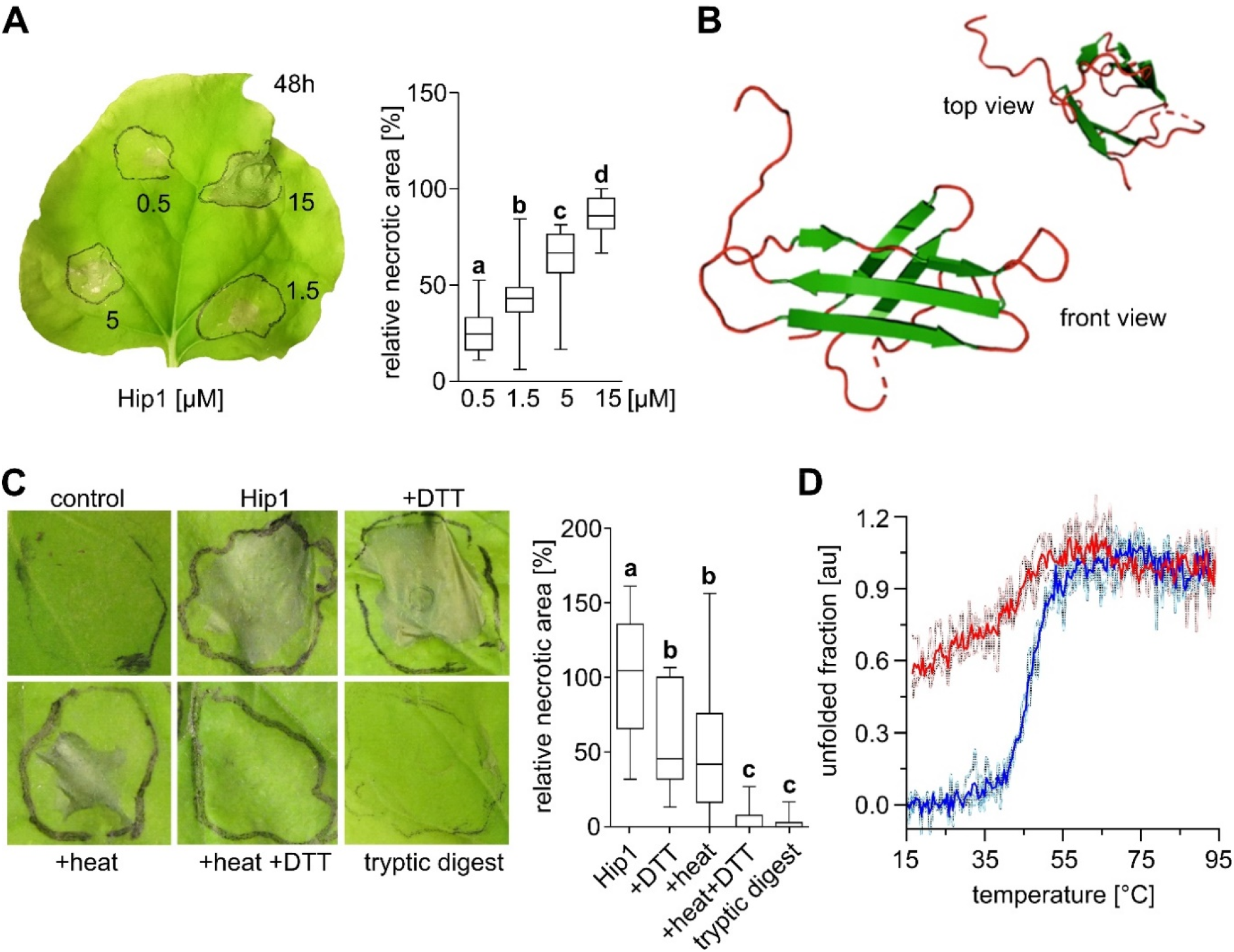
Evidences of the role of tertiary structure in Hip1 activity. Necrotic activity of different Hip1 protein concentrations was analysed by protein infiltration into *N. benthamiana*. One-way ANOVA followed by Dunnett’s multiple comparison post hoc test. *b: p* <0.01; c: *p* <0.001, d: *p* <0.001; n=20 **(A)**. 3D protein structure prediction using Phyre 2 (http://www.sbg.bio.ic.ac.uk/~phyre2/) **(B)**. Effects of different treatments of Hip1 protein on its phytotoxic activity One-way ANOVA followed by Dunnett’s multiple comparison post hoc test. *p-value:* different letters represent *p* <0.01; n>12 **(C)**. Thermal un- (blue) and refolding (red) of Hip1 monitored by circular dichroism (CD) spectroscopy **(D)**.

Given that most of the Hip1 sequence is required for toxicity, 3D structure prediction was carried out using the Phyre2 server, which performs modelling of sequences based on template sequences with known structure (Kelley et al., 2015) (Fig. 3B). Based on X-ray crystallography of the *Alternaria alternata* allergen Alt 1 a (Chruszcz et al., 2012), a ß-barrel conformation was predicted with 80% confidence (Fig. S4C-D). Notably, antiparallel ß-sheets, forming a barrel-like structure are shared by MoHrip1 from *Magnaporthe oryzae* (Zhang et al., 2017) and Pevd1 from *Verticillium dahliae* (Zhang et al., 2019) (Fig. S4D), and thus are considered as members of the family of Alt a 1 proteins. In order to investigate whether this putative fold is required for mediating toxicity, we challenged the folded state of Hip1 by different means. Dithiothreitol (DTT) was used to reduce disulfide bonds, heat treatment employed for protein denaturation and tryptic digest to hydrolyse peptide bonds. This resulted in a significant reduction of necrosis-inducing activity in case of DTT and heat treatment alone but around 50% activity remained whereas the combination of DTT and heat treatment as well as tryptic digest almost entirely abolished toxicity (Fig. 3C). To understand why toxicity was only partially decreased upon heat treatment, we tested Hip1 heat stability by circular dichroism (CD) spectroscopy. For this purpose, a temperature gradient ranging from 15°C to 95°C was applied, and the CD signal measured. As expected, this revealed a temperature-induced conformational change of Hip1 (Fig. 3D, blue curves) which, however, was partly reversed upon cooling (Fig. 3D, red curves). Although refolding was only partial, the observation that the midpoint of the temperature downscan fell close to that of the upscan indicates that part of the protein regained its native conformation even after exposure to high temperature.

As recognition of PAMPs by PRRs usually takes place at the plasma membrane in plants (Boutrot and Zipfel, 2017), we aimed to confirm Hip1 localization by using live-cell imaging. A Hip1-GFP fusion displayed similar levels of necrosis-inducing activity (Fig. S5A and B) and, when transiently expressed, was found at cell borders of *N. benthamiana* leaf epidermal cells (Fig. 4A and B). Colocalization studies, using the plasma membrane marker spRFP-TMD23 (Scheuring et al., 2012) showed Hip1-GFP in close proximity to the PM (Fig. 4C). Furthermore, the Hip1 signal seemed often to be localized between neighbouring epidermal cells, indicating its position in the apoplast (Fig. 4D). To test whether Hip1 indeed functions as a PAMP, typical immune reactions indicative for a PTI were assessed. To this end, reactive oxygen species (ROS) formation and ethylene accumulation were measured in *N. benthamiana* leaves upon Hip1 treatment. For both experimental setups the classical bacterial flagellin-derived PAMP-elicitor flg22 (Felix et al., 1999) was used as positive control. Indeed, Hip1 treatment led to the accumulation of ROS (Fig. 4E) and ethylene (Fig. 4F). Together, this strongly suggests PAMP activity of Hip1. Notably, no significant ethylene accumulation was observed in *Arabidopsis*, and also only weak necrosis was induced by Hip1 (Fig. S5C-E).

**Fig. 4:**
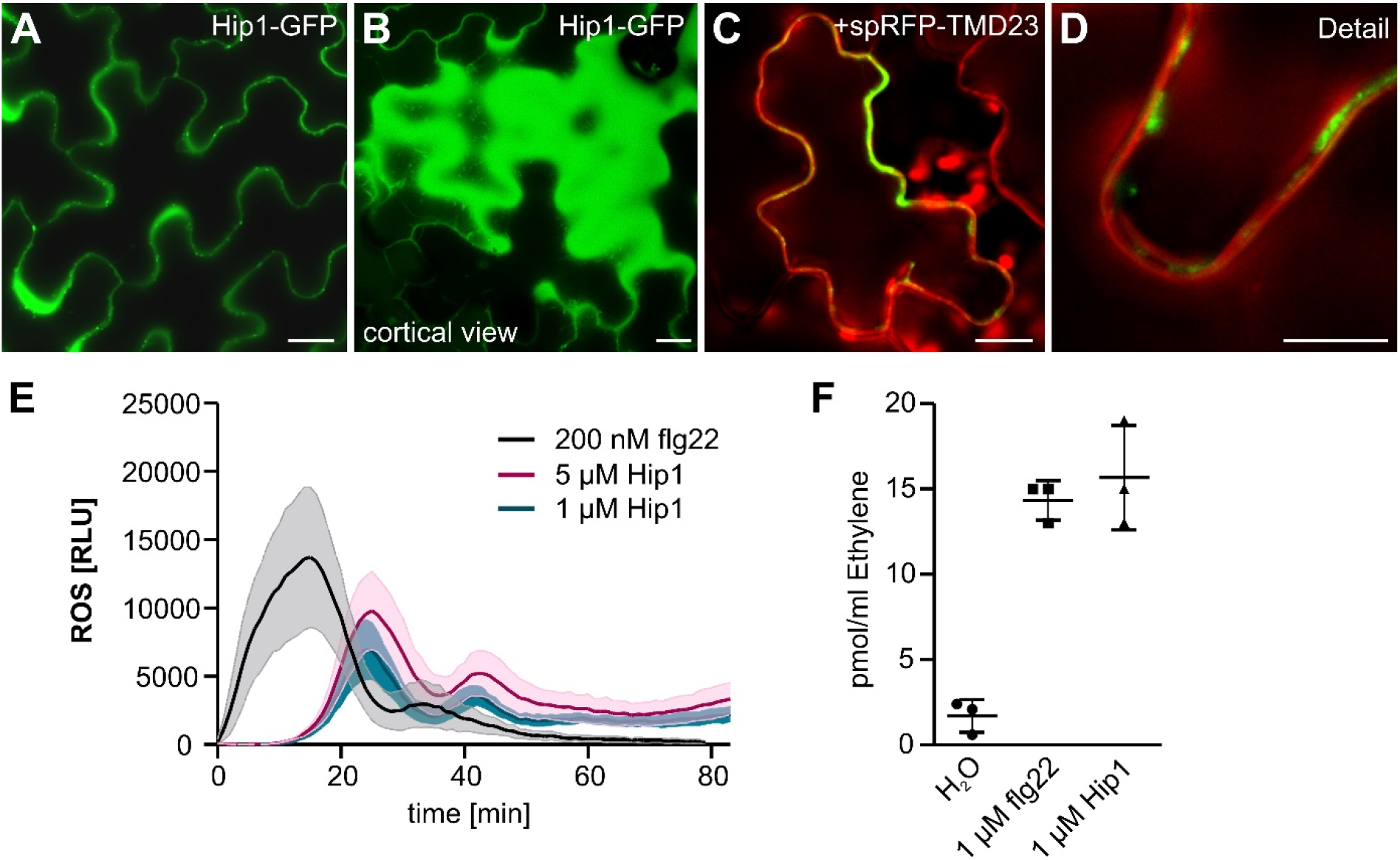
Hip1 is secreted and recognized as pathogen-associated molecular pattern (PAMP). Hip1-GFP highlights the borders of epidermal *N. benthamiana* cells when transiently expressed **(A).** The cortical view of transformed cells indicate secretion of Hip1-GFP into the apoplast **(B)**. Co-expression of Hip1-GFP with the plasma membrane marker spRFP-TMD23 **(C** and **D)**. Scale bars: 15 μm. Reactive oxygen species (ROS) measurement of different Hip1 protein concentrations and the prototypical PAMP flg22 as control **(E)**. n= 4 leaf discs; experiment was repeated 3 times with similar results. Error bars represent standard error of the mean Ethylene accumulation 3h after treatment with Hip1 and flg22 as control for PAMP activity **(F)**. Each data point represents 3 leaf discs measured in one vial. The experiment was repeated 3 times with similar results.

## Discussion

The infection success of *Botrytis* has been reported to depend on the secretion of a complex mixture of cell wall degrading enzymes, necrosis-inducing proteins, phytotoxic metabolites, small RNAs and various organic acids (van Kan, 2006; Weiberg et al., 2013; Veloso and van Kan, 2018; Müller et al., 2018). However, impairment of the proteinaceous portion of the secretome largely abolishes necrosis-inducing activity (Fig. S1; (Zhu et al., 2017)). This suggests that it is mainly proteins that are contributing to toxicity and questions a major role of other metabolites and compounds. By fractionating the *on planta* secretome and analyzing candidate proteins for their phytoxicity by agroinfiltration, we here identified a previously unknown necrosis-inducing protein – Hip1 (Fig. 1). Although several toxic proteins have been identified over the last two decades in *Botrytis*, for many of them it remains elusive how they induce host cell death. Interestingly, the secreted proteins Xyn11A (Noda et al., 2010) and IEB1 (Frías et al., 2016) are able to trigger necrosis induction independently of enzymatic function. For both, it has been shown that short peptides (30 aa and 35 aa) are sufficient to induce PTI and this in turn seems to be linked to their toxic activity. The observed partial inactivation of Hip1 by heat and DTT, separately (Fig. 3), could be explained by partial refolding upon cooling and limited access of cysteine residues within the folded protein. Irreversible unfolding, however, largely abolished the necrosis-inducing toxicity of Hip1, indicating a crucial role of the tertiary structure of the protein. Heat treatments also do not abolish the activity of several other necrosis-inducing proteins, like PG3 (Zhang et al., 2014b) and SsCut from *Sclerotinia sclerotiorum* (Zhang et al., 2014a), and in case of the latter the entire C-terminus has been demonstrated to be dispensable for defense induction. Conversely, the tertiary structures of XYG1 (Zhu et al., 2017) and Spl1 (Frías et al., 2014; Frías et al., 2016) have been shown to be crucial for necrosis and PTI induction, as heat treatment indeed abolishes their function. In case of Hip1, no short peptide conferring toxicity could be identified (Fig. 2 and S4). If Hip1 is recognized by a PAMP receptor, this could indicate that recognition depends on an epitope that is formed only upon proper protein folding. Localization at the plasma membrane and the induction of typical immune responses suggests that Hip1 toxicity indeed relies on its recognition as PAMP (Fig. 4). Notably, 3D protein structure prediction strongly suggests that Hip1 belongs to a fungal class of structural, but not sequence-related proteins, termed the AltA1 family (Pfam family PF16541, (Chruszcz et al., 2012)). For two members of this family, defense and necrosis-inducing activity has been reported (Wang et al., 2012; Zhang et al., 2017; Zhang et al., 2019) but their biological function is still unclear.

Although, PTI is usually associated with basal immunity, many PAMP-like proteins have been shown to induce cell death during the plant HR, typically a feature of effector triggered immunity (ETI) (Jones and Dangl, 2006). For instance, CBEL, a protein from *Phytophthora parasitica var nicotianae*, is recognized as PAMP and induces HR-like lesions and defense responses (Gaulin et al., 2002; Khatib et al., 2004). Certain harpins from *Pseudomonas syringae* likewise represent PAMPs that seem to interfere with host membrane integrity (He et al., 1993; Engelhardt et al., 2009). In this context, it seems plausible that the necrosis-inducing activity of Hip1 results from HR-associated cell death. Indeed, plant cell death has contrasting roles in defense against necrotrophs and biotrophs. While the HR in response to biotrophic pathogens is an indicator of resistance, for several necrotrophic pathogens the HR is even beneficial (Mengiste, 2012; Pitsili et al., 2020). Enhancing plant cell death has been demonstrated to increase susceptibility against *Botrytis* and might even be considered as hallmark of successful infection by necrotrophs (Govrin and Levine, 2000; Govrin et al., 2006; Veronese et al., 2004). Several *Botrytis* CWDEs and NIPs possess PAMP activity in addition to their enzymatic or pore-forming function. Even PAMP activity of proteins without enzymatic function (e.g. IEB1 and Spl1) has been shown to be sufficient to induce necrosis concomitant with plant defense reactions. Therefore, knockout of individual proteins from the toxic cocktail of secreted proteins might not reveal a phenotype but collectively together they could be causal for the broad host spectrum of *B. cinerea*.

Interestingly, both Hip1-induced ROS accumulation and toxicity in *Arabidopsis* was considerably weaker than in *N. benthamiana* (Fig. S5). This could indicate that induction of plant defence and toxic activity are tightly linked and not separable as shown for other secreted toxic proteins, e.g. XYG1 (Zhu et al., 2017). If this holds true, the only known biological function of Hip1 is the activation of plant immune responses to induce enhanced plant cell death. This mechanism has been described for necrotrophic effectors but their necrotic activity is restricted to host plants with an appropriate genotype (Oliver and Solomon, 2010). Since Hip1 homologs are widespread in plant pathogenic and plant-associated ascomycetous fungi (Table S2), it seems plausible that toxicity is mediated via non-canonical PAMP-activity. The restricted toxicity in *Arabidopsis* suggests furthermore that the putative Hip1 receptor is missing here. Recently, a proteinaceous cell death-inducing PAMP from pathogenic fungi has been described as being exclusively active in Solanaceae including *N. benthamiana* belongs (Franco-Orozco et al., 2017). The cell death inducing activity of another PAMP, VmE02 from the necrotrophic fungus *Valsa mali*, has been shown to depend on the presence of a specific receptor like protein (RLP) at the PM of *N. benthamiana* (Nie et al., 2019; Nie et al., 2020). It is conceivable that the differential Hip1 toxicity is relying on binding of a Solanaceae-specific, yet unknown, receptor at the cell surface.

## Material and Methods

### Plant materials and growth conditions

For leaf infiltration and transient expression, *N. benthamiana* was grown for 4-6 weeks on soil in long-day conditions (14h light/10h dark) at 23°C. *Arabidopsis thaliana* ecotype Col-0 grown under long-day regime (16h light/8h dark) for 4 weeks at 22°C was used for infiltration and ethylene accumulation experiments.

### Fungi

*Botrytis cinerea* B05.10 was used as wild type strain for infection tests, growth tests, secretome production and as genetic background for the Hip1 *knockout* generation.

### Recombinant plasmid construction

Coding sequences were amplified from *Botrytis cinerea* cDNA or plasmid DNA. Recipient vectors and fragments were cut using indicated restriction sites (Table S3). Vectors were dephosphorylated prior ligation and fragment purified via gel-extraction if necessary. For transient expression, pGreen II derivatives (pFF04 and pFF18) were used as recipient vectors (Künzl et al., 2016). Both vectors harbour a cauliflower mosaic virus (CMV) 35S promotor, an N-terminal signal peptide (SP) and a kanamycin resistance cassette. A hemagglutinin (HA)-tag for immunological detection was added at the C-terminus of the coding sequence via the reverse primers. The Hip1-GFP fusion was constructed by GreenGate cloning (Lampropoulos et al., 2013), using the following modules: 35S promotor (pGGA004), ER signal sequence (pGGB006), linker-GFP (pGGD001) and RBCS terminator (pGGE001) The Hip1 coding sequence devoid of *BsaI* restriction sites for GreenGate cloning was synthesized by Biocat (Germany). For recombinant protein expression, the Hip1 sequence was amplified without signal peptide and cloned into the pET28a (+) expression vector (Addgene, USA), eventually having a His-tag at the N-terminus. All primers used are shown in Table S3. Gene constructs were verified by sequencing (Seq-It, Germany). For colocalization studies, the PM marker spRFP-TMD23 (pFK44) was used (Scheuring et al., 2012).

### RNA extraction and quantitative real-time PCR

Total RNA of *Botrytis* infected leaves was extracted using the NucleoSpin RNA Plant kit (Macherey-Nagel, Germany). RNA samples were reverse transcribed using the iScript cDNA synthesis kit (Bio-Rad, USA). qPCR was performed with the PerfeCTa SYBR Green SuperMix (Quantabio, USA) in a CFX Connect Real-Time PCR Detection System (Bio-Rad, USA).

Target specific and control primers are listed in Table S3. Expression levels were normalized to the expression levels of Actin 2. Relative expression ratios were calculated according to the efficiency corrected calculation model (Pfaffl, 2001).

### Cultivation and transformation of *Botrytis*

Cultivation of *Botrytis* was performed as described by (Müller et al., 2018).

### Secretome isolation and proteomics

Secretome samples were isolated from tomato leaves 48h after *Botrytis* inoculation, as described previously (Müller et al., 2018).

### Transient gene expression

*Agrobacterium tumefaciens* (strain GV3101::mp90) was electroporated in the presence of 500 ng binary vector (pGreen II) using a gene pulser II (Biorad, USA) set to 2.5 kV, 25 μF and 400 Ω. For transient expression, overnight grown liquid cultures of transformed agrobacteria were washed with infiltration medium (Sparkes et al., 2006) and adjusted to OD 0.3 prior infiltration in *N. benthamiana*. Subsequently, the infiltrated area was marked and assessed after 72h incubation. The relative necrotic area was calculated as ratio of the infiltrated area and the necrotic area. For co-expression, agrobacteria cultures were adjusted to OD 0.15 and mixed before infiltration.

### Recombinant protein expression

*E.coli* strains Rosetta pLysS (Novagen, USA) and T7 Shuffle (New England Biolabs, USA) were used to express Hip1. Protein expression was induced by 0.4 mM IPTG and conducted at 37°C for 3 h or at 20°C over night. After bacterial lysis, protein purification was carried out by immobilized metal affinity chromatography (IMAC) using His/Ni beads (Roth, Germany) as resin. Subsequently, dialysis was carried out using either 25 mM citrate buffer (pH5) or 50 mM phosphate buffer with 150 mM NaCl (pH6). For ethylene and ROS measurements, Hip1 protein was dialysed against deionized H_2_O.

### Protein extraction and immunological detection

Proteins were extracted from leaf disks (20 mm diameter) of infiltrated tobacco by grinding in liquid nitrogen using mortar and pestle. The frozen powder was suspended in extraction buffer (100 mM potassium, 20 mM HEPES, 0.1 mM EDTA, 2 mM DTT, 1 mM PMSF; pH 7.5) and after centrifugation protein determination by Bradford was carried out. Equal protein amounts were separated via SDS-PAGE and subsequently subjected to immunoblot on nitrocellulose membrane. Monoclonal antibodies against hemagglutinin coupled to peroxidase (Roche; Cat. No. 12013819001, 1:1000 dilution) or GFP (Roche;11814460001, 1:500 dilution) were used for chemiluminescence detection (ECL Prime Kit; Amersham, UK) on a FUSION detection system (Vilber Lourmat, FR).

### Confocal microscopy

Images were acquired using a Zeiss LSM880 AxioObserver confocal laser scanning microscope equipped with a Zeiss C-Apochromat 40x/1,2 W AutoCorr M27 water-immersion objective (DFG, INST 248/254-1). Fluorescent signals of GFP (excitation/emission 488 nm/500-571 nm) and RFP (excitation/emission 543nm/580-718 nm) were processed using the Zeiss software ZEN 2.3 or ImageJ (https://imagej.nih.gov/ij/). For co-expression studies, images were acquired using sequential scan mode to avoid channel crosstalk.

### Circular dichroism spectroscopy

Thermal un- and refolding was monitored between 15°C and 95°C at 215 nm with a Chirascan-plus spectrometer (Applied Photophysics, UK) equipped with a 150-W xenon arc lamp. The heating/cooling rate was 1°C/min. Samples contained about 15 μM Hip1 and were measured using a 1 mm quartz glass cuvette (Hellma, Germany). Traces were normalized by assuming a completely folded state for the first 15 data points and a completely unfolded state for the last 5 data points. The mean of three experiments is shown in bold, individual measurements are displayed on light blue/red.

### Ethylene measurements

*Nicotiana benthamiana* leaf discs (3 mm) floated overnight on ddH_2_O. Three leaf discs were put into one glass vial (6ml volume) with 200 μl ddH_2_O. The respective elicitor or ddH_2_O as control were added and the vials were closed with a rubber cap. The samples were shaken for 3h at 100 rpm on a horizontal shaker. 1ml air was taken out through the rubber cap with a syringe and injected into a gas chromatograph. Each treatment was measured with 3 replicates.

### ROS assays

Reactive oxygen species (ROS) production was measured as described in (Ranf et al., 2015). Small leaf discs (3 mm diameter) of *Nicotiana benthamiana* plants were cut and floated overnight on 200 μl ddH_2_O in a 96-well plate (one disc per well; white 96-well plates). Before starting the measurement, the water was removed with a pipette and 75 μl of Luminol-Mastermix (2 μg/ml horseradish peroxidase (Type II, Roche), 5μM L-012 (WAKO chemicals)) was added into each well. Luminescence was measured as relative light units (RLU) in 1min intervals using a Tecan F200 luminometer. The background was measured for 10min. The elicitors were always added in a volume of 25 μl diluted with Luminol-Mastermix. After elicitor treatment luminescence readings were continued for 90min. Data is shown after normalization to average ROS levels 5 min before elicitor application and subtraction of water controls that were included for each genotype on the same plate.

### Statistical analysis

Analysis was carried out using the GraphPad Prism software. The detailed statistical method employed is provided in the respective figure legends. All experiments were carried out at least three times. Box limits in the graphs represent 25th to 75th percentile, the horizontal line the median and whiskers minimum to maximum values.

## Acknowledgements

We would like to thank Peter Pimpl for providing the vectors pFF04 and pFF18. Furthermore, technical help from Oliver Trentmann, Sabrina Kaiser, Tabea Lang, Florian Fässler, Marco Aras and Benjamin Ledermann is highly appreciated. For the kind willingness to share facilities and equipment, we thank Michael Schroda and Nicole Frankenberg-Dinkel. We are grateful to Amir Sharon for critical reading of the manuscript.

## Author’s contribution

CS, MH, and DS designed research. TJ, TL, CS, JM, and DS conducted most experiments; FS performed MS-MS analysis; FM and SK carried out CD spectroscopy. TJ, TL, CS, JM, FM, MH, and DS analysed data. DS created the figures and wrote the manuscript. All co-authors commented on the manuscript.

