## Supplemental Information for "The secreted hypersensitive response inducing protein 1 from *Botrytis cinerea* displays non-canonical PAMP-activity"

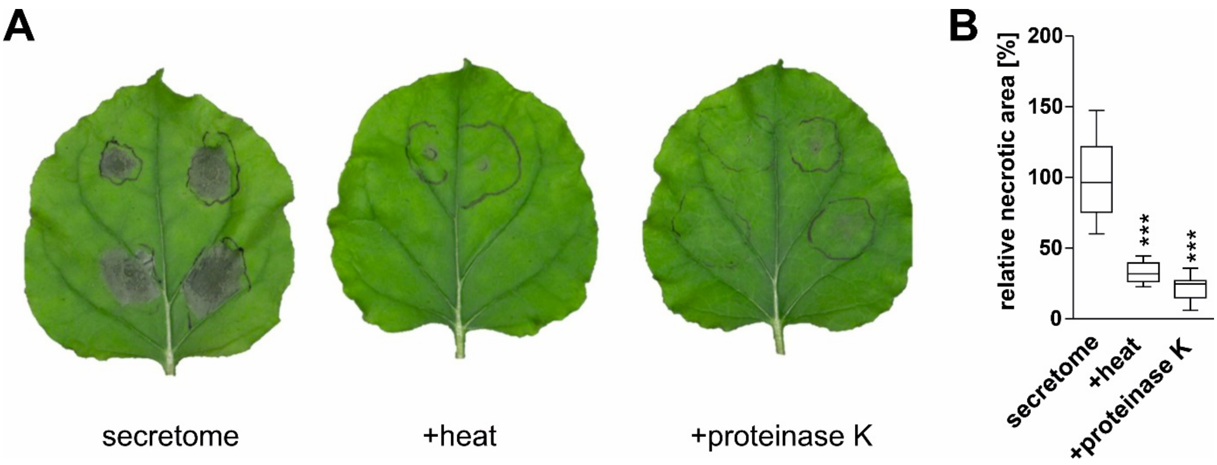

**Figure S1: Toxicity of secreted proteins and other compounds from *Botrytis* rely on protein activity.** Isolation of secreted proteins and substances 48h after *Botrytis* infection of *Solanum lycoperiscum* and subsequent infiltration into *N. benthamiana* leaves **(A)**. Heat and proteinase K treatment reduced the secretome´s toxicity **(B)**. One-way ANOVA followed by Tukey´s post hoc test with untreated secretome as control. *p-value*: *** p<0.001; n=9 (secretome), n=5 (+heat), n=7 (+proteinase K).

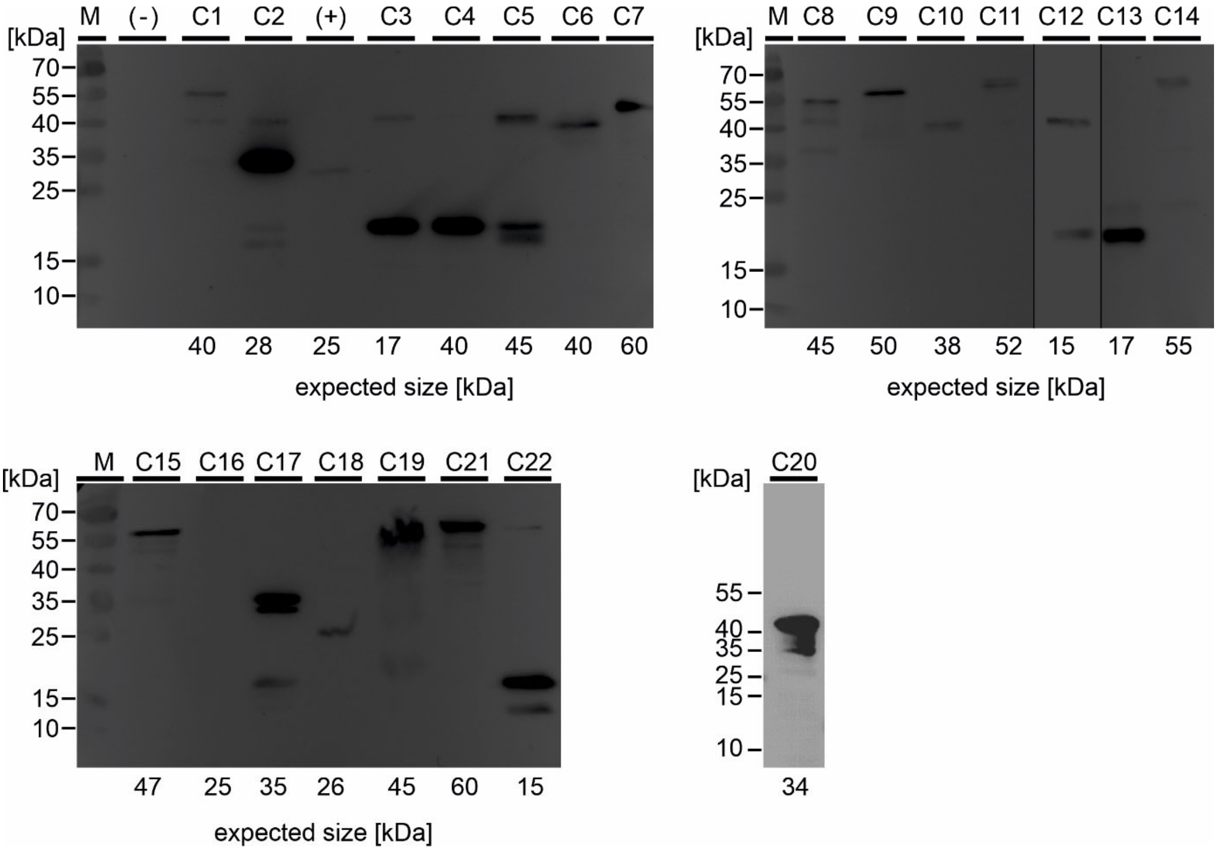

**Figure S2: Expression control of candidates from *Botrytis*.** Proteins were extracted 3 dpi and separated via SDS-PAGE. After blotting, an HA-antibody coupled to peroxidase was used for visualization of signals.

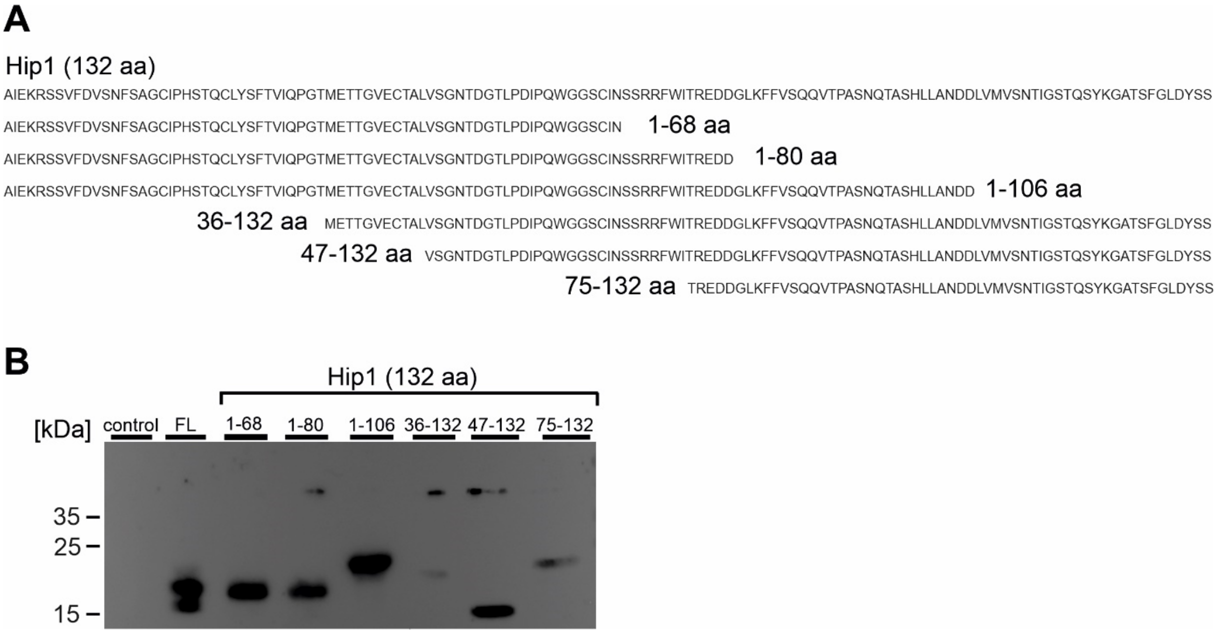

**Figure S3: Expression control of truncated Hip1 derivatives.** Overview of tested constructs **(A)**. Westernblot of Hip1 derivatives using an HA-antibody coupled to peroxidase **(B)**.

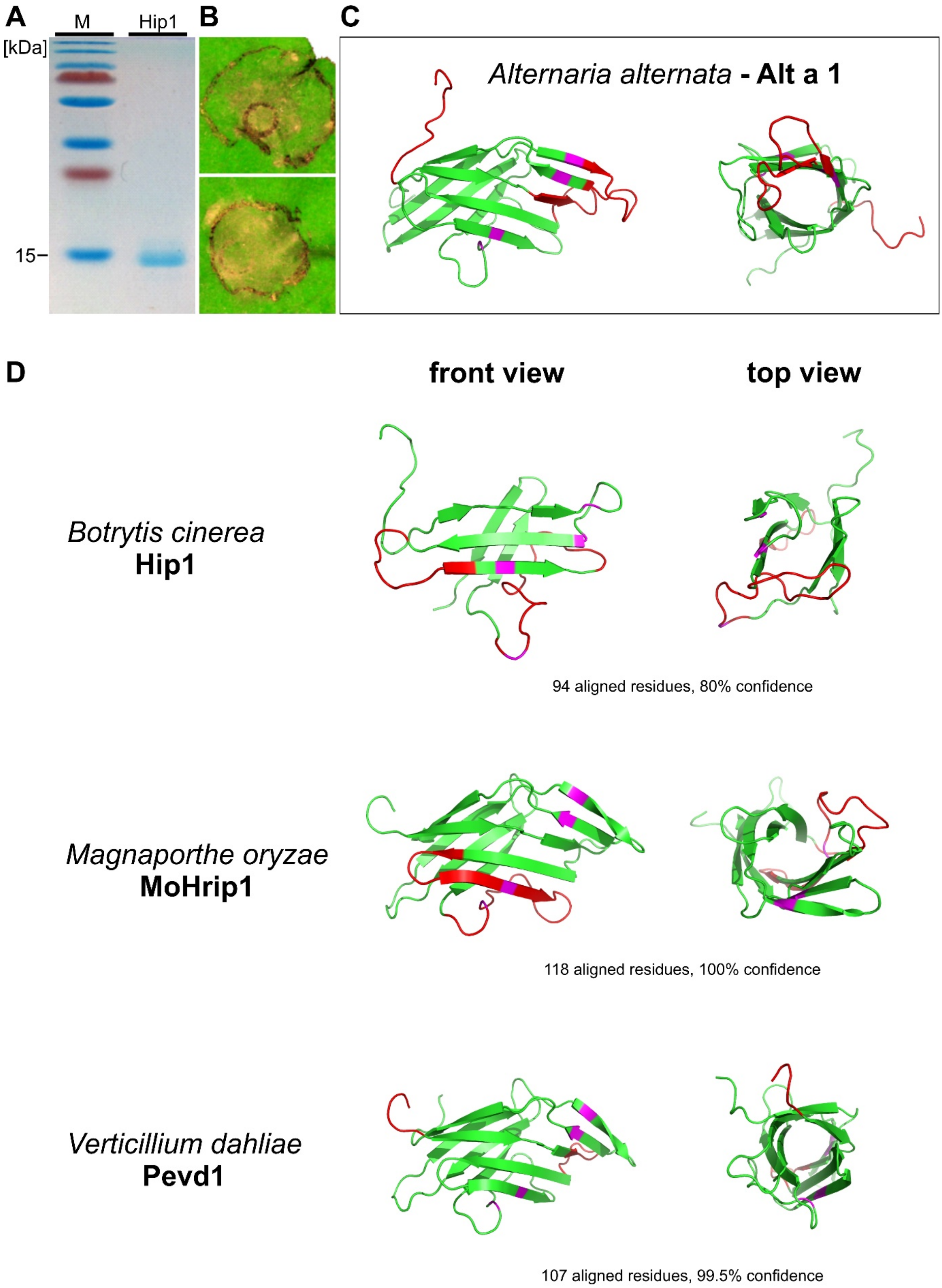

**Figure S4: Protein expression and structure prediction of Hip1.** The *E.coli* Rosetta (DE3) host strain was used to express Hip1 upon IPTG induction. After purification, a single band, predicted at ~15 kDa was visible **(A)**. At lower concentrations (5 µM), infiltrated Hip1 protein caused necrotic spots which resemble sites of plant hypersensitive response **(B)**. Based on x-ray crystallography of the *Alternaria alternate* allergen Alt 1 a **(C)**, Hip1 tertiary structure was predicted using the software Phyre2 (http://www.sbg.bio.ic.ac.uk/~phyre2/). Typical antiparallel ß-sheets forming a barrel-like structure are features shared by other fungal proteins as well **(D)**. Coverage of the original amino acid sequence and confidence of prediction are indicated below the 3D models. Cysteins are highlighted in pink.

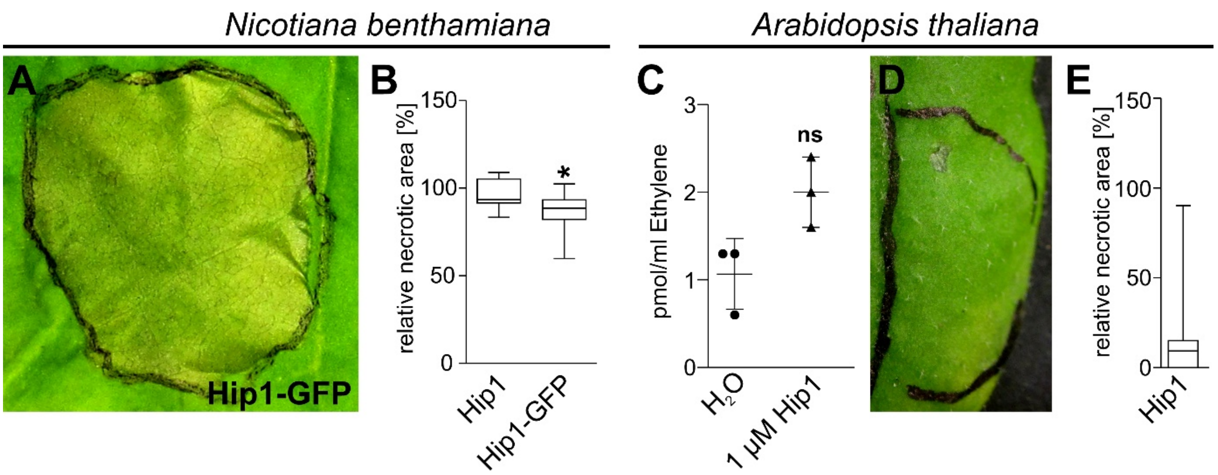

**Figure S5: Hip1 induces strong necrosis in *N. benthamiana* but not in *Arabidopsis*.** In transient expression, Hip1 and Hip1-GFP induces full necrosis in tobacco after 72h (**A** and **B**). Ethylene measurements upon Hip1 treatment in *Arabidopsis* Col-0 wt **(C)**. Infiltration of 15 µM Hip1 (48h, n=20) did not induce clear necrosis in the *Arabidopsis* (**D** and **E**). Student t-test; *p-value*: *$p<0.05$; ns= not significant.

#### Table S1

| Candidate No | Gene Accession | Function [PFAM output/assigned] | Fold change [in vitro vs infection] | Size [kDa] |
|---|---|---|---|---|
| C1 | Bcin01g06010 | *Glycosyl hydrolases family 16* | -2.1 | 40 |
| C2 | Bcin02g07100 | *GDSL-like Lipase / Acylhydrolase family* | 12.8 | 28 |
| C3 | Bcin03g05720 | *unknown* | 30.5 | 17 |
| C4 | Bcin06g06410 | *Fungal cellulose binding domain* | 165 | 40 |
| C5 | Bcin07g02650 | *Cellulase (glycosyl hydrolase family 5)* | 215 | 45 |
| C6 | Bcin14g00850 | *Glycosyl hydrolases family 28 /* ***BcPG1*** | 3.1 | 40 |
| C7 | Bcin15g03150 | *Pro-kumamolisin, activation domain /* ***Sedolisin*** | 3.5 | 60 |
| C8 | Bcin01g11160 | *Glycosyl hydrolases family 28 /* ***Exopolygalacturonase*** | 17.1 | 45 |
| C9 | Bcin08g00280 | *Serine carboxypeptidase* | 12 | 50 |
| C10 | Bcin08g02970 | *Pectinesterase /* ***BcPME1*** | 2.4 | 38 |
| C11 | Bcin09g04790 | *Glycosyl hydrolases family 39* | 47 | 52 |
| C12 | Bcin11g06400 | *unknown* | -2 | 15 |
| C13 | Bcin13g00300 | *unknown* | 6 | 17 |
| C14 | Bcin13g03950 | *Alpha-L-arabinofuranosidase B (ABFB)* | 293 | 55 |
| C15 | Bcin14g03970 | *Glucanosyltransferase* | -3 | 47 |
| C16 | Bcin14g04980 | *SNARE* | -1.5 | 25 |
| C17 | Bcin14g05510 | *Glycosyl hydrolase family 62* | 30,5 | 35 |
| C18 | Bcin15g00130 | ***Cutinase*** | 330 | 26 |
| C19 | Bcin16g03950 | *Glycosyl hydrolase family 7* | 1739 | 45 |
| C20 | Bcin01g02460 | *Rare lipoprotein A (RlpA)-like double-psi beta-barrel /* ***Expansin-like*** | 2 | 24 |
| C21 | Bcin11g01310 | *TAP-like protein* | -1.6 | 60 |
| C22 | Bcin14g01200 | *unknown /* ***HR-inducing protein*** | -1.5 | 15 |

#### Table S2

```
Botrytis cinerea     MQFSSAIISAITVALASAAAIEKRSSVFDVSNFSAGCIPHSTQCLYSFTVIQPGTMETTG
O. ulmi              MKFSAAAILSAVVAVSANPTVARAEPVFAVSNFQAGCIPHSSQCRYSFDVIKTGTGETIP
F. graminearum       MKFSAAILAA------AAATGATADAVLNVRDFAASCTPHSAMCSYSFNVIQPGTMDTKG
C. higginsianum      MKFAAVLS---SAAVAAAAAIGKRDVTFSVSDFSAGCIPHSTQCLIAFNVIQPGTMETTG
                     *:*::.          :  :    . .: * :* *.* ***: *  :* **: ** :*

Botrytis cinerea     VECTALVSGNTDGTLPDIPQWGGSCINSSRRFWITREDDGLKFFVSQQVTPASNQTASHL
O. ulmi              VSCVLLKT--SNNVLPDVTD--GTCIDSSRTFSFKRRAEGLTFTVSQQITPSSNQTGTYL
F. graminearum       YECTAKLPGAGPGELPEVKV--GTCLPSSRTFDVVRSDKGLTLTVSVQVSPNSFTKGSHL
C. higginsianum      VECRALVPAKSDGTLPDVKE--AACTESSRTFDLVRSPEGITFTVSQPVTPSSNQTGSHL
                      .*         . **::    .:*  *** * . *  .*:.: **  ::* *  ..::*

Botrytis cinerea      LANDDLVMV--SNTIGSTQSYKGATSFGLDYSS-
O. ulmi               IPKNDFIFT--KTTTASVEEYTGPKAFTLTDQTI
F. graminearum        IPTKDIKTVKGDTPTGDAQAYVGPKDFPLERVD-
C. higginsianum       LPSSEFVIS--NQPNAVVESYTGPNAFDLE----
                      : ..::     .   . .: * * . * *
```

### Table S3

| **Candidate Screening** | | | | | |
|---|---|---|---|---|---|
| **Name** | **Accession** | **Primer Forward** | **Sequence** | **Primer Reverse** | **Sequence** |
| XYG1 | Bcin03g03630 | Bcin03g03630_FW_SalI | TAGGCAGTCGACTAACCCTACTCCTACTCTTGAG | Bcin03g03630_HA_RV_SpeI | ATCTACTAGTTTAAGCGTAATCTGGAACATCGTATGGGTAATTGAGCGAGACGGAGTA |
| Candidate 1 | Bcin01g06010 | Bcin01g06010_FW_NheI | TAGGCAGCTAGCGCCCAAACATTCACTGATTGCAAC | Bcin01g06010_HA_RV_BamHI | ATCTGGATCCTTAAGCGTAATCTGGAACATCGTATGGGTACATAACGAGATAACCTAATCC |
| Candidate 2 | Bcin02g07100 | Bcin02g07100_FW_NheI | TAGGCAGCTAGCGCCGTGCCAACTTTGTATTTGGCT | Bcin02g07100_HA_RV_BamHI | ATCTGGATCCCTAAGCGTAATCTGGAACATCGTATGGGTAAGCACAGCTTCCGGC |
| Candidate 3 | Bcin03g05720 | Bcin03g05720_FW_NheI | TAGGCAGCTAGCGCCGCTCCAACTGCAATCCCA | Bcin03g05720_HA_RV_BamHI | ATCTGGATCCCTAAGCGTAATCTGGAACATCGTATGGGTATGCCTCGCAAGCATA |
| Candidate 4 | Bcin06g06410 | Bcin06g06410_FW_SalI | TAGGCAGTCGACTCAGCAGGCCGTGTAT | Bcin06g06410_HA_RV_SpeI | ATCTACTAGTTTAAGCGTAATCTGGAACATCGTATGGGTACCAAAATACACCCTTTAAGAGT |
| Candidate 5 | Bcin07g02650 | Bcin07g02650_FW_NheI | TAGGCAGCTAGCGCCTCTCCAATTGAAGAACGAGCC | Bcin07g02650_HA_RV_BamHI | ATCTGGATCCTTAAGCGTAATCTGGAACATCGTATGGGTATGGCGAGTAAATCTTCGT |
| Candidate 6 | Bcin14g00850 | Bcin14g00850_FW_SalI | TAGGCAGTCGACTGCTCCAGCACCAGCA | Bcin14g00850_HA_RV_SpeI | ATCTACTAGTTTAAGCGTAATCTGGAACATCGTATGGGTAACACTTGACACCAGATGG |
| Candidate 7 | Bcin15g03150 | Bcin15g03150_FW_NheI | TAGGCAGCTAGCGCCTTGCCCACTTTCGGCAAA | Bcin15g03150_HA_RV_XbaI | ATCTTCTAGATTAAGCGTAATCTGGAACATCGTATGGGTAAGTAGAATTACTGATTGCCAACAA |
| Candidate 8 | Bcin01g11160 | Bcin01g11160_FW_NcoI | TAGGCACCATGGGATCACCAACTGCAGTGAAAAGC | Bcin01g11160_HA_RV_BamHI | ATCTGGATCCTTAAGCGTAATCTGGAACATCGTATGGGTATCCCGACGCACCAGA |
| Candidate 9 | Bcin08g00280 | Bcin08g00280 FW NcoI | TAGGCACCATGGGAGCCCCTACCACTGAAGGG | Bcin08g00280_HA_RV_BamHI | ATCTGGATCCCTAAGCGTAATCTGGAACATCGTATGGGTAAGTAGAAGAGAGAGGCTTCTTGC |
| Candidate 10 | Bcin08g02970 | Bcin08g02970_FW_SalI | TAGGCAGTCGACTGCTCCTCTAGAGGCACGA | Bcin08g02970_HA_RV_SpeI | ATCTACTAGTTTAAGCGTAATCTGGAACATCGTATGGGTACAAATAACTGCTATCAACCCACCA |
| Candidate 11 | Bcin09g04790 | Bcin09g04790_FW_NheI | TAGGCAGCTAGCGCCTCAAATATAATACAGAGACAAACTATC | Bcin09g04790_HA_RV_BamHI | ATCTGGATCCCTAAGCGTAATCTGGAACATCGTATGGGTAACTGACATTGAATTCAAAAGCA |
| Candidate 12 | Bcin11g06400 | Bcin11g06400_FW_NheI | TAGGCAGCTAGCGCCTCTCCTGTCCCCG | Bcin11g06400_HA_RV_BamHI | ATCTGGATCCTTAAGCGTAATCTGGAACATCGTATGGGTAGCATTTAAATCGTTCAGATAGAAG |
| Candidate 13 | Bcin13g00300 | Bcin13g00300_FW_SalI | TAGGCAGTCGACTAGCACATCCGACGCT | Bcin13g00300_HA_RV_SpeI | ATCTACTAGTCTAAGCGTAATCTGGAACATCGTATGGGTATTGAGCAATCAATTGGATCTGCTC |
| Candidate 14 | Bcin13g03950 | Bcin13g03950_FW_NcoI | TAGGCACCATGgcGGGACCTTGCGATATCTACTC | Bcin13g03950_HA_RV_BamHI | ATCTGGATCCttaAGCGTAATCTGGAACATCGTATGGGTAAGCAAAGCCGGTGC |
| Candidate 15 | Bcin14g03970 | Bcin14g03970_FW_NcoI | TAGGCACCATGGccGCTCCATCTCCAGTTGAGA | Bcin14g03970_HA_RV_XbaI | ATCTTCTAGattAAGCGTAATCTGGAACATCGTATGGGTACAAAAGTGCAGCACCA |
| Candidate 16 | Bcin14g04980 | Bcin14g04980_FW_SalI | TAGGCAGTCGACGgAGCTCAAGCTCGTCC | Bcin14g04980_HA_RV_SpeI | ATCTACTAGTttaAGCGTAATCTGGAACATCGTATGGGTATCTGAACTTGCTAACAATAACAG |
| Candidate 17 | Bcin14g05510 | Bcin14g05510_FW_NheI | TAGGCAGCTAGCGCTTGTGCTCTCCCATC | Bcin14g05510_HA_RV_BamHI | ATCTGGATCCttaAGCGTAATCTGGAACATCGTATGGGTACTTCTTGAGAGTCAAAACTCC |
| Candidate 18 | Bcin15g00130 | Bcin15g00130_FW_NcoI | TAGGCACCATGgcGCTCCCTTCTGAATTGGACAC | Bcin15g00130_HA_RV_XbaI | ATCTTCTAGattAAGCGTAATCTGGAACATCGTATGGGTAACCAGATGAAGCGAGC |
| Candidate 19 | Bcin16g03950 | Bcin16g03950_FW_Nhe I | TAGGCAGCTAGCgcaCAACAAGTTGGTACCTATCAAAC | Bcin16g03950_HA_RV_BamHI | ATCTGGATCCttaAGCGTAATCTGGAACATCGTATGGGTAACCGTAAGTAGAGTTAATAGCTC |
| Candidate 20 | Bcin01g02460 | Bcin01g02460_FW_SalI | TAGGCAGTCGACTGCCGTGATGGGAAAG | Bcin01g02460_HA_RV_SpeI | ATCTACTAGTCTAAGCGTAATCTGGAACATCGTATGGGTAGGAAGTCTCGCACTC |
| Candidate 21 | Bcin11g01310 | Bcin11g01310_FW_SalI | TAGGCAGTCGACTCTGACATGGACCTCATGT | Bcin11g01310_HA_RV_SpeI | ATCTACTAGTTTAAGCGTAATCTGGAACATCGTATGGGTATGCGATTGCTAAGAAGGC |
| Candidate 22 | Bcin14g01200 | Bcin14g01200_FW_SalI | TAGGCAGTCGACTGCCATCGAGAAGCGC | Bcin14g01200_HA_RV_SpeI | ATCTACTAGTCTAAGCGTAATCTGGAACATCGTATGGGTAGGAGCTGTAGTCCAAGCC |
| **Hrp1 truncations** | | | | | |
| Hrp1 1-68 | | Bcin14g01200_FW_SalI | TAGGCAGTCGACTGCCATCGAGAAGCGC | Hrp1_N-terminus no2_SpeI_RV | ATCTACTAGTCTAAGCGTAATCTGGAACATCGTATGGGTAGTTGATGCAGGATCCTCCCCA |
| Hrp1 1-80 | | Bcin14g01200_FW_SalI | TAGGCAGTCGACTGCCATCGAGAAGCGC | Hrp1_REDD_RV | ATCTACTAGTCTAAGCGTAATCTGGAACATCGTATGGGTAGTCGTCCTCGCGGGTGAT |
| Hrp1 1-106 | | Bcin14g01200_FW_SalI | TAGGCAGTCGACTGCCATCGAGAAGCGC | Hrp1_LANDD_RV | ATCTACTAGTCTAAGCGTAATCTGGAACATCGTATGGGTAGTCGTCGTTGGCGAGCAAG |
| Hrp1 36-132 | | Hrp1_METT_FW | TAGTCAGTCGACTATGGAGACCACCGGTGTAGAG | Bcin14g01200_HA_RV_SpeI | ATCTACTAGTCTAAGCGTAATCTGGAACATCGTATGGGTAGGAGCTGTAGTCCAAGCC |
| Hrp1 47-132 | | Hrp1_C-terminus_SalI_FW | TAGGCAGTCGACGGTCTCCGGCAACACCGAC | Bcin14g01200_HA_RV_SpeI | ATCTACTAGTCTAAGCGTAATCTGGAACATCGTATGGGTAGGAGCTGTAGTCCAAGCC |
| Hrp1 75-132 | | Hrp1_C-terminus no2_SalI_FW | TAGGCAGTCGACTCGCGAGGACGACGGTC | Bcin14g01200_HA_RV_SpeI | ATCTACTAGTCTAAGCGTAATCTGGAACATCGTATGGGTAGGAGCTGTAGTCCAAGCC |
| **Recombinant Hrp1 expression** | | | | | |
| Bcin14g01200_FW_NdeI | | CAACATATGGCCATCGAGAAGCGCAGCAG | | | |
| Bcin14g01200_RV_EcoRI | | TGTGAATTCCTAGGAGCTGTAGTCCAAGCCGAA | | | |
| **Greengate cloning of Hrp1-GFP** | | | | | |
| GG_fulllength-hrp1_FW | | AACAGGTCTCAGGCTCAACAATGCAATTCTCATCTGCCATTATCTCTGC | | | |
| GG_fulllength-hrp1_REV | | AACAGGTCTCTCTGAGGAGCTGTAGTCCAAGCCGAAG | | | |
| ***Botrytis* CRISPR knockout Hrp1** | | | | | |
| Hrp1 gRNA1 | aagcTAATACGACTCACTATAgCTCCCAGACATCCCACAATGgTTTTAGAGCTAGAAATAGCAAG | | | | |
| Hrp1 gRNA2 | aagcTAATACGACTCACTATAGTTGGCGAGCAAGTGGGAAGgTTTTAGAGCTAGAAATAGCAAG | | | | |
| Hrp1 RT FW | GCAAAAATGCAATTCTCATCTGCCATTATCTCTGCCATCACCGTTGCTCTTGCATCGGCCTGCTGGCCTTTTGCTCACATGCATG | | | | |
| Hrp1 RT RV | GTGCAGACATTCGCAGCAAGCGAAAGCCATGACGCTTTGCCACCTCGGTCGATTCCTCTAATCGCCGGAAAGGACCCGCAAATG | | | | |
